# Disulfide cross-linked redox-sensitive peptide condensates are efficient cell delivery vehicles of molecular cargo

**DOI:** 10.1101/2025.05.20.655132

**Authors:** Malay Mondal, Windfield S. Swetman, Shazeed-Ul Karim, Sabin Shrestha, Ashe M. Davis, Fengwei Bai, Faqing Huang, Tristan D. Clemons, Vijayaraghavan Rangachari

**Author notes:** Correspondence to Vijay Rangachari.

## Abstract

Biomolecular condensates (BCs) are phase-separated viscoelastic hubs within demixed solutions enriched in proteins and nucleic acids. Such condensates, also called membraneless organelles, are increasingly observed in cells and serve as transient hubs for spatial organization and compartmentalization of biomolecules. Along with the transiency of formation and dissolution, their ability to sequester molecules has inspired us to develop BCs as potential vehicles to transport and deliver molecular cargo. We recently reported the design of disulfide bond cross-linked phase-separating peptide (PSP) condensates that spontaneously dissolve in reducing conditions (*JACS, 2024, <u>146</u>, 255299*). Based on the premise that the highly reducing cytoplasm could dissolve PSP condensates and release partitioned cargo, here, we demonstrate the ability of PSP condensates to deliver molecular cargo to the cytoplasm of HeLa cells efficiently. We show that PSP condensates deliver a variety of cargos that differ in their sizes and chemistries, including small molecules, peptides, GFP protein (31 kDa), DNA (1.7 kbp), and mRNA. The transfection efficiencies of PSP condensates for delivering DNA and mRNA were also significantly greater than those of a commercial transfection agent. With room to tailor the condensate properties based on cargo and cell types, these results showcase the potential of disulfide-cross-linked PSPs as effective and customizable cellular delivery vehicles, filling a critical demand gap for such delivery systems.

## INTRODUCTION

Efficient cell delivery is key to many therapeutic and pharmaceutical applications. The dearth of effective cellular delivery systems was exacerbated by the COVID-19 pandemic, which brought to the fore some glaring issues with the delivery of mRNA cargo in cells^1,2^. Standard delivery systems such as lipid nanoparticles (LNP), viral vectors, and nanocarriers suffer instability, inefficiency, immunogenicity, and toxicity, in addition to manufacturing difficulty and long-term safety^3^, severely blunting their effectiveness and utility^4–9^. Synthetic polymer hydrogels have emerged as potential alternatives but are largely incompatible for physiological use^10,11^. In pursuing alternative and better cell delivery solutions, protein-based delivery vehicles have increasingly become attractive due to their compatibility with cells and the ease of customizability toward tailoring physical, chemical, and material properties^11–13^. In particular, biomolecular condensates (BCs) have emerged as promising soft materials for molecular delivery, transport, and tissue engineering due to their viscoelastic properties ^13–16^.

In cells, BCs are dense foci enriched in proteins and nucleic acids formed transiently by demixing from homogenous solutions^17–23^. Devoid of a lipid membrane coat, these BCs are also commonly known as membraneless organelles, which play many pathological roles in cells^24–26^. BCs are formed when proteins undergo a density transition accompanied by phase separation, commonly liquid-liquid phase separation (LLPS), by which they demix from the bulk solution to generate at least two immiscible phases: a protein-rich dense phase and a protein-depleted dilute phase^24,25,27–29^. Multivalent transient interactions involving cation-π and π-π between arginine and tyrosine (Arg-Tyr) and Tyr-Tyr residues respectively, underlie the molecular forces within BCs ^24,25,27–29^. The sequence determinants governing the formation of BCs are well captured by a ‘stickers and spacers’ model in which multivalent interactions by the stickers, such as Arg and Tyr, and volume confinement by the disorder-promoting spacers, such as Gly, Ala, and Ser residues, drive phase separation^17,28^. Phase separation occurs above a threshold called the saturation concentration (*C_sat_*)^28,30^. Increasing the concentrations further will enter another regime called a percolation threshold (*C_perc_*) that enables network-spanning interactions that drive coacervates into viscoelastic fluids^25,29,30^. BCs can be formed from homotypic interactions called ‘self-coacervation’ or heterotypic interactions with other molecules called ‘complex-coacervation,’^17^ the latter being prevalent in cells.

In our previous study, we designed phase-separating peptides (PSPs) containing ‘sticker and spacer’ sequences and interspersed them with cysteines (Cys) in specific locations^31^. We showed that in oxidizing conditions, PSPs not only formed stable BCs but maintained their morphology and viscoelasticity for strikingly prolonged periods, establishing, for the first time, the significance of disulfide covalent cross-links in BCs^31^. We also demonstrated that under reducing conditions, the condensates dissolved completely and that the formation and dissolution of the condensate droplets are reversible under redox flux ^31^. Finally, we showed that molecular cargo can be partitioned within these condensates with encapsulation efficiencies depending on the chemistry of the cargo, with pore sizes of PSPs large enough to accommodate the cargo. Based on these results, we hypothesize that since the extracellular environment is oxidizing and the cytoplasm is reducing^32^, PSP condensates are efficient, safe, and customizable vehicles for transporting and delivering molecular cargo into the cellular cytoplasm.

Here, we tested this hypothesis using HeLa cells and PSP-2 condensates as a model delivery system, with a host of cargo ranging from small molecules (Fluorescein isothiocyanate (FITC), fluorescein, and FITC-PSP-2) to large biopolymers such as DNA encoding enhanced green fluorescence protein (EGFP), mRNA transcript of GFP, and GFP itself, which vastly differ in their chemistries and sizes (Figure 1). With a variety of biophysical and cell biological techniques, we discovered that PSP-2 condensates efficiently encapsulate all the cargo investigated, cross the cell membrane to dissolve, and release cargo in the cytoplasm spontaneously. These results demonstrate the utility of redox-sensitive designer peptide condensates as a novel class of customizable cell delivery vehicles.

**Figure 1.**
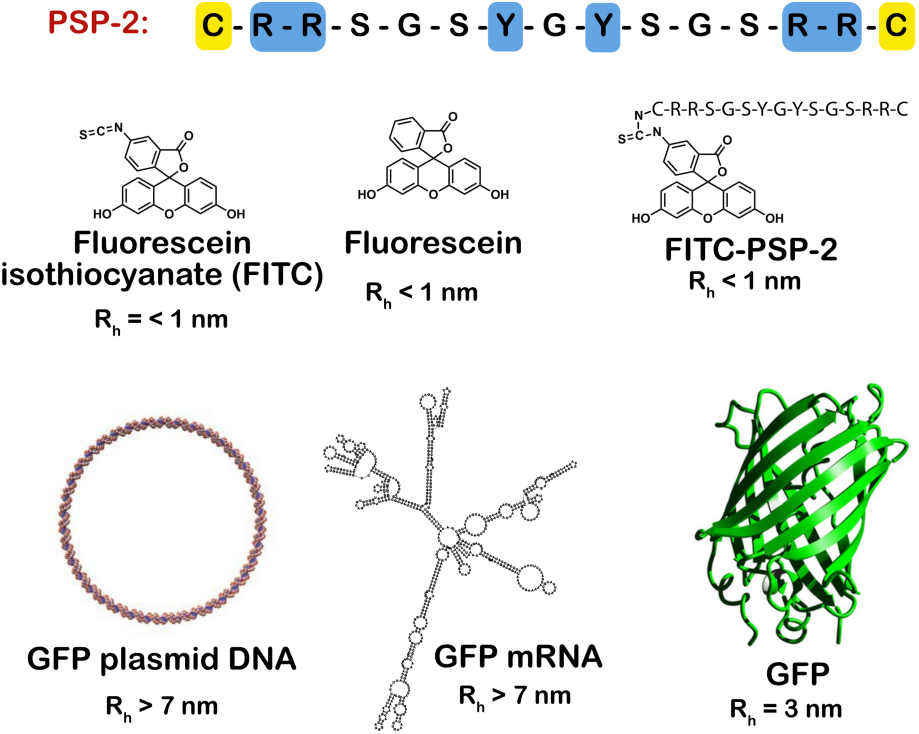
PSP-2 peptide sequence and molecular cargos used in this study. (Top) The primary sequence of the peptide PSP-2 with stickers highlighted in blue and cysteines in yellow. (Bottom) Chemical structures of small molecules, schematic of GFP DNA plasmid, predicted structure of GFP mRNA (RNAfold web server), and the structure of GFP (PDB: 4KW4), which are used as cargo. The approximate predicted hydrodynamic radii (R_h_) of each cargo are shown.

## RESULTS AND DISCUSSION

We first examined the condensate formation of PSP-2 in buffer conditions. Our results reconfirmed the previously established characteristics of PSP-2 condensates^31^. Briefly, PSP-2 formed condensates both by self-coacervation and complex-coacervation with polyA-RNA, as confirmed by confocal and brightfield imaging (Figure S1a-d). Fluorescence recovery after photobleaching (FRAP) analysis demonstrated liquid-like characteristics of the condensates with >90% recovery (Figure S1c and f). Also consistent with our previous finding, the addition of DTT completely dissolved self-coacervates, as observed in confocal microscopic images, and the droplets reformed upon treatment with 2% H_2_O_2_, an oxidizing agent (Figure S1i). The reversibility of condensates was also confirmed by FRAP analysis of the droplet before the addition of the reducing agent and after re-oxidation, both of which showed nearly identical recovery rates (Figure S1j). These data confirm the expected physiochemical behavior, and redox sensitivity of PSP-2 condensates in buffers.

### PSP-2 condensates are stable in cell culture medium and are not cytotoxic to HeLa cells

Before investigating cargo delivery into cells, it is imperative to establish whether the condensates can be stable in cell medium and concentration regimes. More importantly, it is important to see if the condensates are cytotoxic. We previously established the *C_sat_* for PSP-2 self-coacervates to be 2.0 mM and complex-coacervates with polyA-RNA to be 50 μM in 50 mM Tris, 2.5 M NaCl, at pH 8.0^31^ (Figure S1). Since Optimum media contains many osmolytes and salts and differs in ionic strength, we anticipated that the medium would likely affect the phase boundary and *C_sat_* values. Therefore, to ascertain whether PSP-2 condensates are stable in cell media, we diluted preformed PSP-2 self-coacervates and complex-coacervates 10- and 20-fold in Optimum® media to final concentrations of 350 and 175 μM for self- and 20 and 10 μM for complex-coacervates. The confocal microscopy images taken after 4 and 24 hours of incubation at 37 °C showed the presence of condensates for both coacervates (Figure 2a and b). FRAP analysis on these droplets showed a robust ∼80% recovery, indicating that the droplets remained viscous (Figure 2c and d). However, confocal images suggested that condensates of PSP-2-RNA complex coacervates were more numerous than the PSP-2 self-coacervates (Figure 2a and b). This difference suggests that the self-coacervates could be less stable than complex-coacervates in cell culture media. This conjecture is also supported by the size difference between PSP-2 and PSP-2-RNA condensates. The average surface area of 0.1 – 0.3 μm^2^ for PSP-2 droplets was significantly smaller than 1.5 – 1.8 μm^2^ for PSP-2-RNA condensates in both concentrations (Figure 2e and f). At 20 μM, PSP-2-RNA droplets showed a dramatic drop in their surface areas from 3.5 μm^2^ after four hours to 1.8 μm^2^ after 24 hours (Figure 2f). More interestingly, the size distribution is reversed between the condensate sizes in buffer versus Optimum® media; while self-coacervates of PSP-2 are larger in the buffer, they are at least two-fold smaller in the media. This is possibly due to many counter-ions interacting with RNA and facilitating their partitioning within the condensates, resulting in larger droplet formation. Nevertheless, the data confirm that both self- and complex- coacervates of PSP-2 can also form in the cell media.

**Figure 2.**
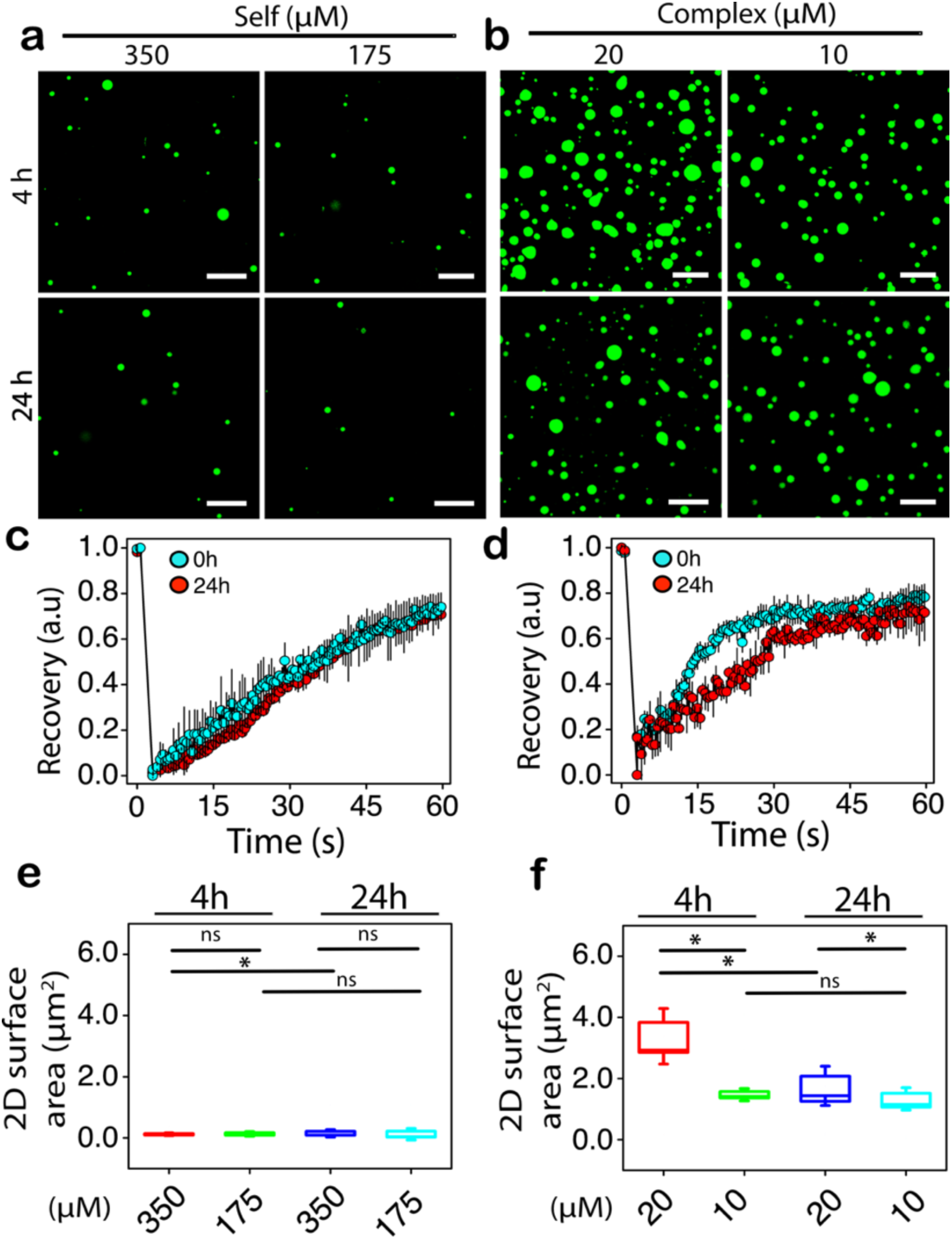
Stability of PSP-2 condensates in cell culture media. (a) Confocal images of PSP-2 self-coacervates (3.5 mM in 50 mM Tris, 2.5 M NaCl, pH 8.0 at 37 °C) containing 1% FITC-labeled PSP-2 diluted to 350 and 175 µM concentrations in Optimum media and imaged after 4 and 24 hours of incubation. (b) Confocal images of PSP-2 complex-coacervates of PSP-2-polyARNA (200 µM peptide and 200 µg/mL RNA in the same buffer but with 150 mL NaCl) diluted to 20 and 10 µM concentrations. (c and d) FRAP data for self- and complex-coacervates in Optimum media. The scale bar is 20 µm. (e and f) Sizes of the droplets were quantified based on the 2D surface area observed in the images using ImageJ software for self-(e) and complex-coacervates (f) after 4 and 24 hours. *\*p* < 0.05; *ns* – not significant based on 2-way ANOVA analysis.

Next, we investigated the potential cytotoxicity of PSP-2 on HeLa cells using a lactate dehydrogenase (LDH) assay measuring LDH release after 4 and 24 hours of incubation. This assay was performed per the manufacturer’s protocol (see Methods). PSP-2 peptides were introduced as phase-separated condensates (self-coacervates; 3.5 mM in 50 mM Tris, 2.5 M NaCl, pH 8.0) and in non-phase-separating buffer conditions (same concentrations without NaCl) at two diluted concentrations (350 µM and 175 µM) in Optimum® media. After four hours of incubation, the results revealed no significant increase in LDH release compared to untreated control cells, showing high cell viability, which suggests that PSP-2 condensates do not compromise cell membrane integrity (Figure 3a). In contrast, the positive control (Triton X-100) induced substantial LDH release, showing less than 10% viability (Figure 3a). Nearly identical results were obtained after 24 hours of incubation (Figure 3b). The absence of significant cytotoxicity within the tested concentration range suggests that PSP-2 condensates maintain cell viability, underscoring their potential for therapeutic delivery applications.

**Figure 3:**
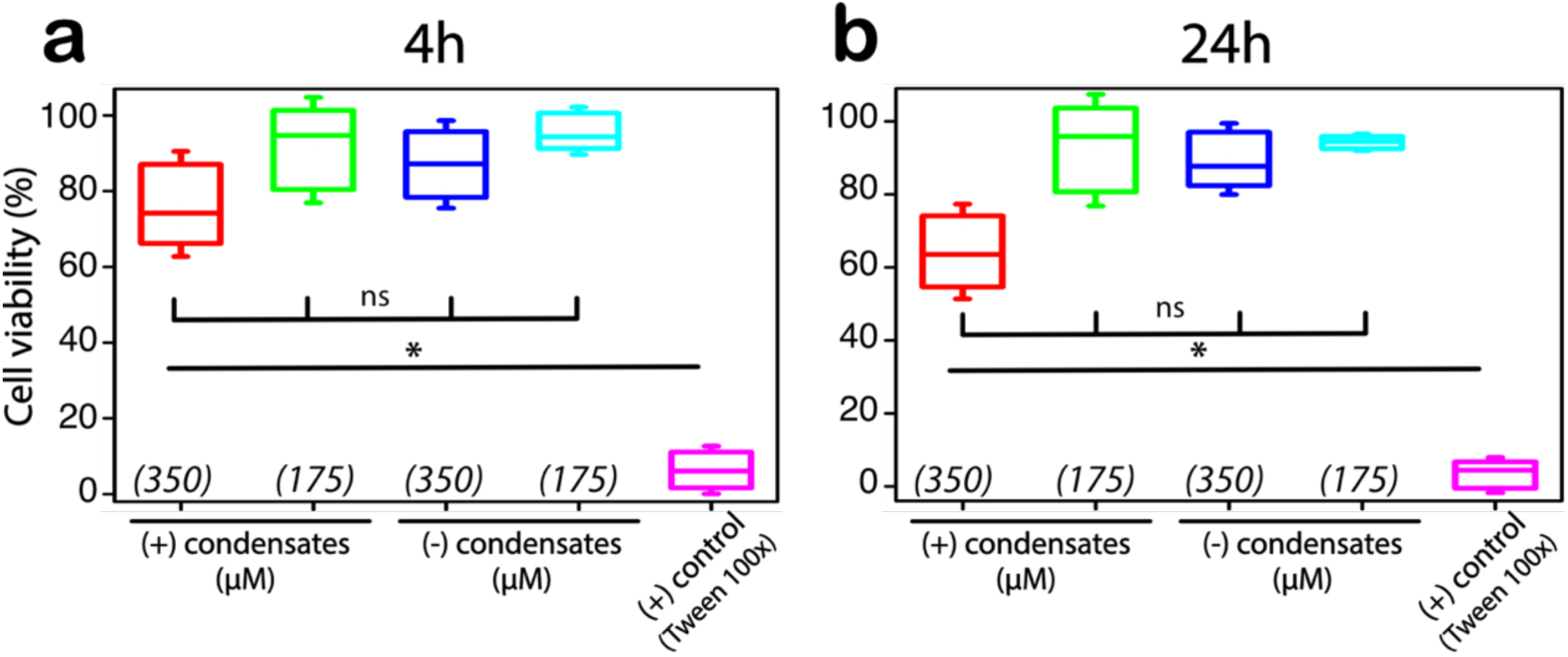
PSP-2 condensates are not toxic to the cells in optimal concentration regimes. Cell viability of HeLa cells in the presence of PSP-2 peptides as condensates (self-coacervates from Figure 1) and in non-phase-separating buffer conditions (3.5 mM in 50 mM Tris, pH 8.0 at 37 °C) diluted to 350 and 175 µM in Optimum media was tested using an LDH assay after 4 hours (a) and 24 h (b) of incubation. Triton X-100 was used as a positive control. All measurements were performed in triplicates. * p<0.05; ns – not significant based on two-way ANOVA.

### PSP-2 condensates efficiently deliver cargo to the cytosol

To investigate the ability of PSP- 2 condensates to deliver cargo intracellularly, we investigated their capacity to transport molecular cargo into HeLa cells by choosing cargos with varying sizes and chemistries. First, FITC and fluorescein are small fluorescent molecules; while FITC forms covalent adducts with proteins through free amine groups, fluorescein is covalently unreactive but engages in non-covalent interactions. Second, we covalently tagged PSP-2 with FITC to see the fate of the peptide in the cells. Third, we used the 31 kDa enhanced green fluorescent protein (EGFP) as a large cargo. Then, we measured the *C_sat_* of cargo-laden condensates in Optimum® media as we expected a change in the phase boundary. Upon serial dilutions of cargo-laden PSP-2 condensates in the media, we observed a drop in the *C_sat_* values between 10 and 25 µM for all the cargo except EGFP, whose value was between 3 and 10 µM (Figure S2). The average condensate size was slightly higher for EGFP-loaded condensates, perhaps due to the partitioning of a larger molecule in the condensates (protein, Figure S3). We also tested the redox sensitivity of the cargo-laden condensates in media by adding the reducing agents DTT and GSH, which showed that all condensates dissolved in reducing environments as expected (Figure S4). Lastly, we determined the encapsulation efficiency by quantifying the amount of cargo in the condensates, as shown previously (Figure S5)^31^ Both FITC and FITC-labeled peptides showed the highest encapsulation efficiency with > 90%, while fluorescein showed the lowest with ∼20%, consistent with our previous observation^31^, while EGFP showed ∼ 50% (Figure S5).

Before incubating with HeLa cells, we confirmed the encapsulation of all the cargos within PSP-2 condensates in Optimum® media by confocal microscopy, which showed encapsulation of FITC (Figure 4a), fluorescein (Figure 4b), FITC-labeled PSP-2 (Figure 4c), and EGFP (Figure 4d) encapsulation with roughly uniform sizes of the complex condensates.

**Figure 4.**
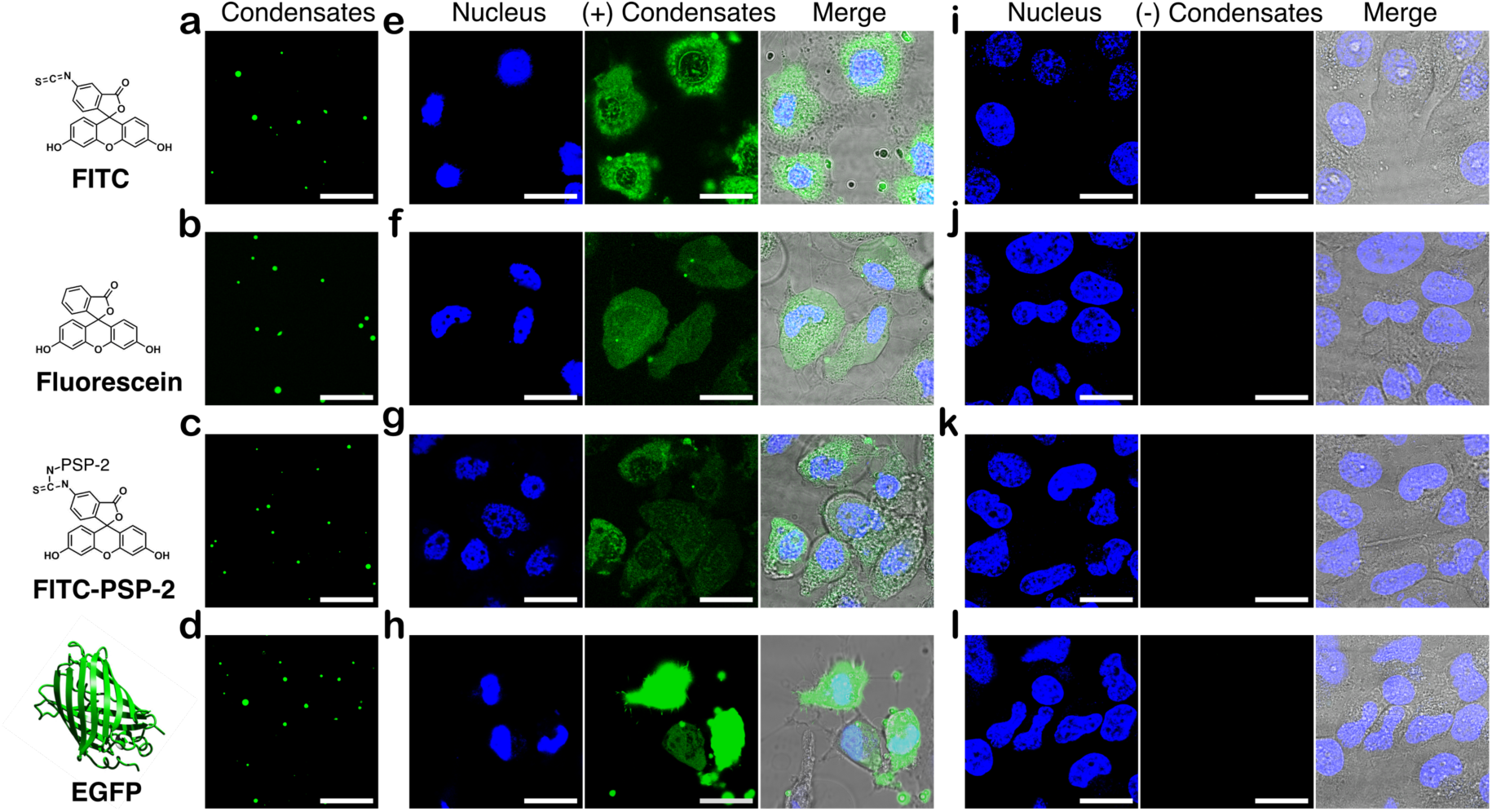
Condensates deliver cargo to the cytosol of HeLa cells. (a-d) Confocal images of cargo-loaded PSP-2 self-coacervates in Optimum media for FITC alone (a), fluorescein (b), FITC-labeled peptides (c), or EGFP protein (d) before passaging to HeLa cells. The cargo concentration with or without condensates was 5 µM, and the condensate concentration was maintained at 350 µM in all the experiments. (e-h) Images of HeLa cells treated with self-coacervates loaded with FITC (d), fluorescein (e), FITC-labeled peptides (f), or EGFP protein (h) were captured after four hours of incubation. (i-l) Confocal images of control HeLa cells after four hours of incubation pre-treated with respective cargo in Optimum media in the absence of condensates. Hoechst33258^®^ (blue) dye was used for nuclear staining. The scale bar is 20 µm.

Following a four-hour incubation with HeLa cells, confocal images revealed robust cytosolic fluorescence signals in condensate-treated cells for all tested cargo molecules: FITC (Figure 4e), fluorescein (Figure 4f), FITC-PSP-2 (Figure 4g), and EGFP (Figure 4h). In contrast, control cells incubated with free cargo without condensates exhibited no fluorescence (Figure 4i-l), indicating ineffective uptake, if any, under identical conditions. These findings establish that PSP-2 condensates transport cargo not only intracellularly but also spontaneously dissolve in the reducing environment of the cytosol, delivering the biomolecular cargo. Future investigations will explore the mechanisms of cargo uptake and potential applications in targeted therapeutic delivery.

Further inspection of the cells after delivery indicates interesting features. Delivery of FITC- laden condensates shows, in addition to diffuse green fluorescence throughout the cytosol indicating cytoplasmic distribution of the cargo upon dissolution of the condensates, some undissolved condensates were also observed (white arrows, Figure 5a). Furthermore, we observed some condensates outside the cells that failed to enter (red arrows; Figure 5a). The 3D reconstruction of the confocal z-stack of the cell viewed from two perpendicular angles demonstrates the distribution of FITC throughout the cytoplasm alongside undissolved condensates (Figure 5b). On the other hand, fluorescein showed a slightly less intense distribution in the cytosol than FITC, likely due to its lower encapsulation efficiency (Figure 5c). We also observed numerous undissolved condensates in addition to the fully dissolved droplets (white arrows; Figure 5c), alongside those that remained outside the cell (red arrows; Figure 5c). This cargo distribution in the cytosol is again observable from the 3D reconstruction of the confocal z-stack (Figure 5d). Similar distributions were also observed for FITC-tagged PSP-2 (Figure 5e and 5f) and EGFP (Figure 5g and 5h), where among all cargo, EGFP shows the most homogenous distribution in the cytosol.

**Figure 5:**
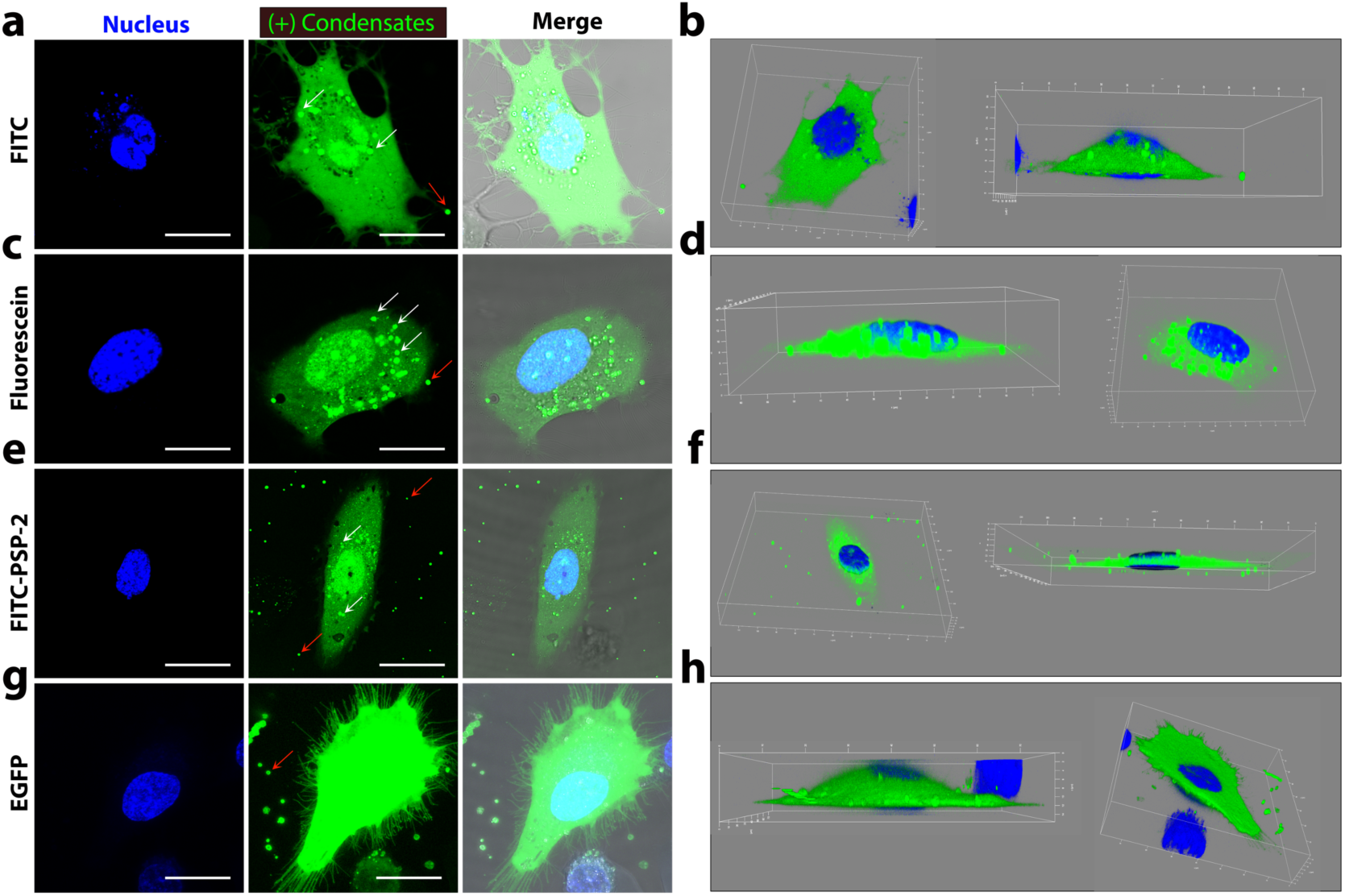
Distribution of condensate-delivered cargo in cellular cytoplasm. Single-cell close-up and 3D images of HeLa cells after the cargo delivery by PSP-2 condensates. (a) Confocal images stained for nucleus (blue) and cargo (green) with merge for FITC. (b) 3D images showing the distribution of FITC. Similar images for cargos, fluorescein (c and d), FITC-PSP-2 (e and f), and EGFP (g and h). White arrows indicate undissolved condensates in the cytoplasm, and red arrows indicate condensates outside the cell. The scale bar is 20 µm.

### PSP-2 condensates efficiently deliver DNA and mRNA to the cytosol

Since the negatively charged polyA-RNA could coacervate with PSP-2, we hypothesized that DNA may also be able to partition into the condensates. To test this hypothesis, we used two types of DNA that encode for EGFP; (*a*) an *E.coli* expressing DNA plasmid containing a T7 promoter with a size of 3589 bp subcloned in a pRSET vector (Figure 6a) and, (*b*) a mammalian cell-expressing linear DNA fragment containing a CMV promoter with a size of 1731 bp (Figure 6b). The CMV-EGFP fragment was generated by PCR using pEG as the template. The DNA plasmid was used to observe the partitioning within PSP-2 condensates, while the linear DNA fragment was used to deliver and express EGFP in HeLa cells. First, to ascertain that the DNA partitioned within PSP-2 condensates, we observed the binding of ethidium bromide (EtBr). EtBr fluorescence was observed under ultraviolet illumination when added to the condensates, which was not observed in samples without condensate formation (Figure 6c). We then centrifuged the DNA-laden condensates and observed the DNA in the sample before centrifugation (T) and the supernatant after (S) by agarose gel electrophoresis (Figure 6d). The gels confirmed that the DNA was partitioned within the condensates as they were not observed in the supernatant (Figure 6d). Encapsulation assessments showed 60% and 85% loading for EGFP plasmid and CMV-EGFP DNA fragment, respectively (Figure 6e). Finally, the DNA’s presence in the condensates was confirmed by confocal microscopy images, which showed the unambiguous presence of DNA using BactoView® stain (Figure 6f and 6g).

**Figure 6:**
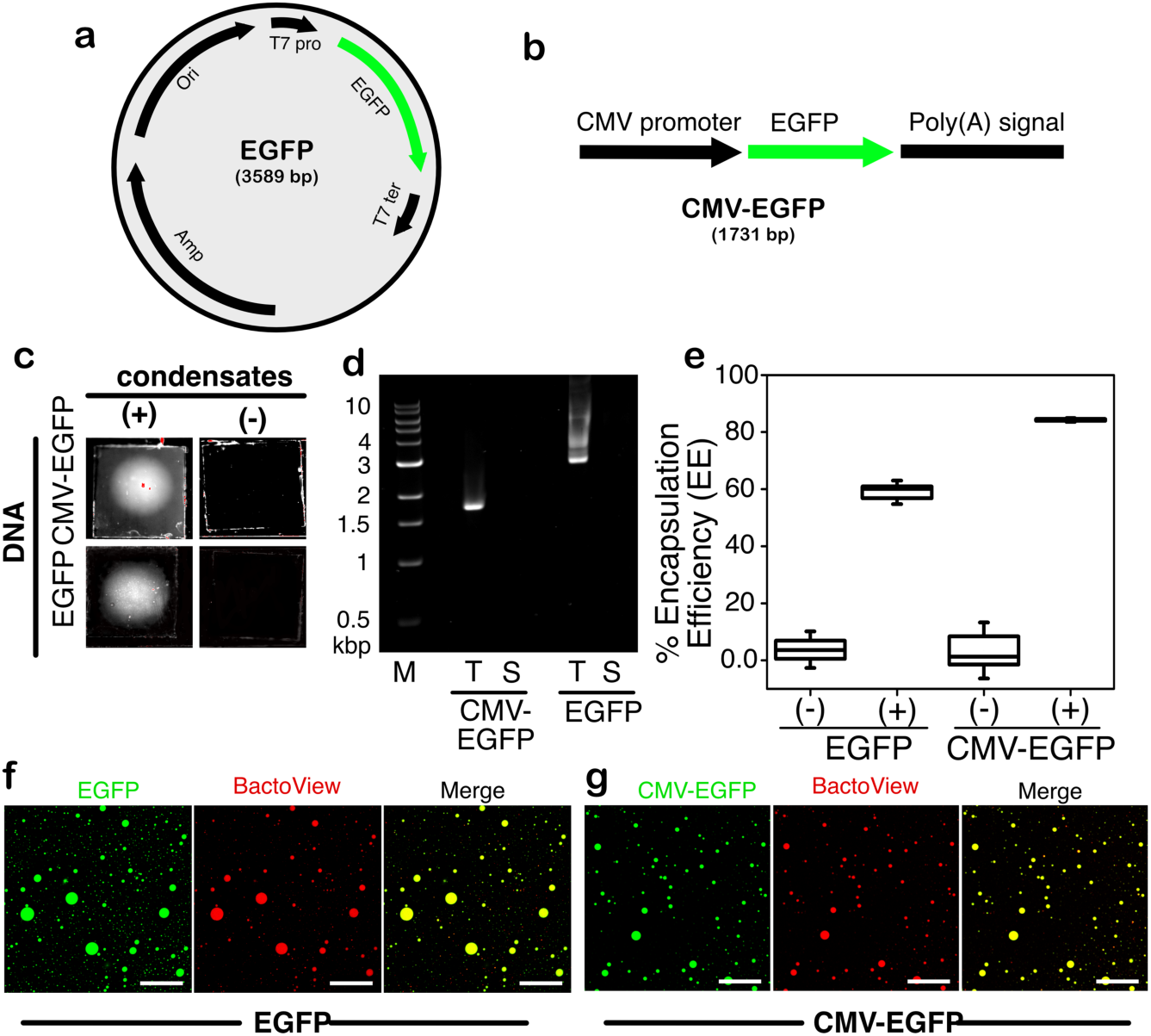
Partitioning of plasmid and linear DNA within PSP-2 condensates. (a-b) Schematic map of EGFP plasmid and the CMV-EGFP linear DNA used in our study. (c) T7-EGFP plasmid and CMV-EGFP DNA fragment with ethidium bromide (EtBr) in the presence and absence of PSP-2 condensates. (d) The DNAs were analyzed using agarose gels (1%) after incubation with condensates. T and S represent the sample before centrifugation and supernatant after centrifugation, respectively. (e) The encapsulation efficiency (EE) of the DNAs within the PSP-2 condensates was calculated as described in Methods; (+) and (-) represent with and without condensates, respectively. (f-g) Confocal microscopy images of DNA-loaded condensates, where the DNAs are stained with BactoView (selective for detecting DNA), and the condensates are visualized using FITC. The scale bar in the images corresponds to 20 µm.

To further investigate the efficiency of a “functional” cargo, we used CMV-EGFP fragment to transfect confluent HeLa cells and compared the transfection efficiency with the commercial Lipofectamine3000®. Upon transfection with CMV-EGFP DNA (1 µg), we observed green fluorescence in the cytoplasm, confirming the expression of EGFP within 6 hours of incubation (Figure 7a). The intensity of EGFP fluorescence increased after 24 and 48 hours, confirming robust expression of EGFP (Figure 7b and 7c). On the other hand, transfection of the same amount of DNA using Lipofectamine300® following the manufacturer’s protocols demonstrated minimal expression levels after 6 hours (Figure 7d), which only marginally improved based on fluorescence intensity after 24 and 48 hours (Figure 7e and 7f).

**Figure 7.**
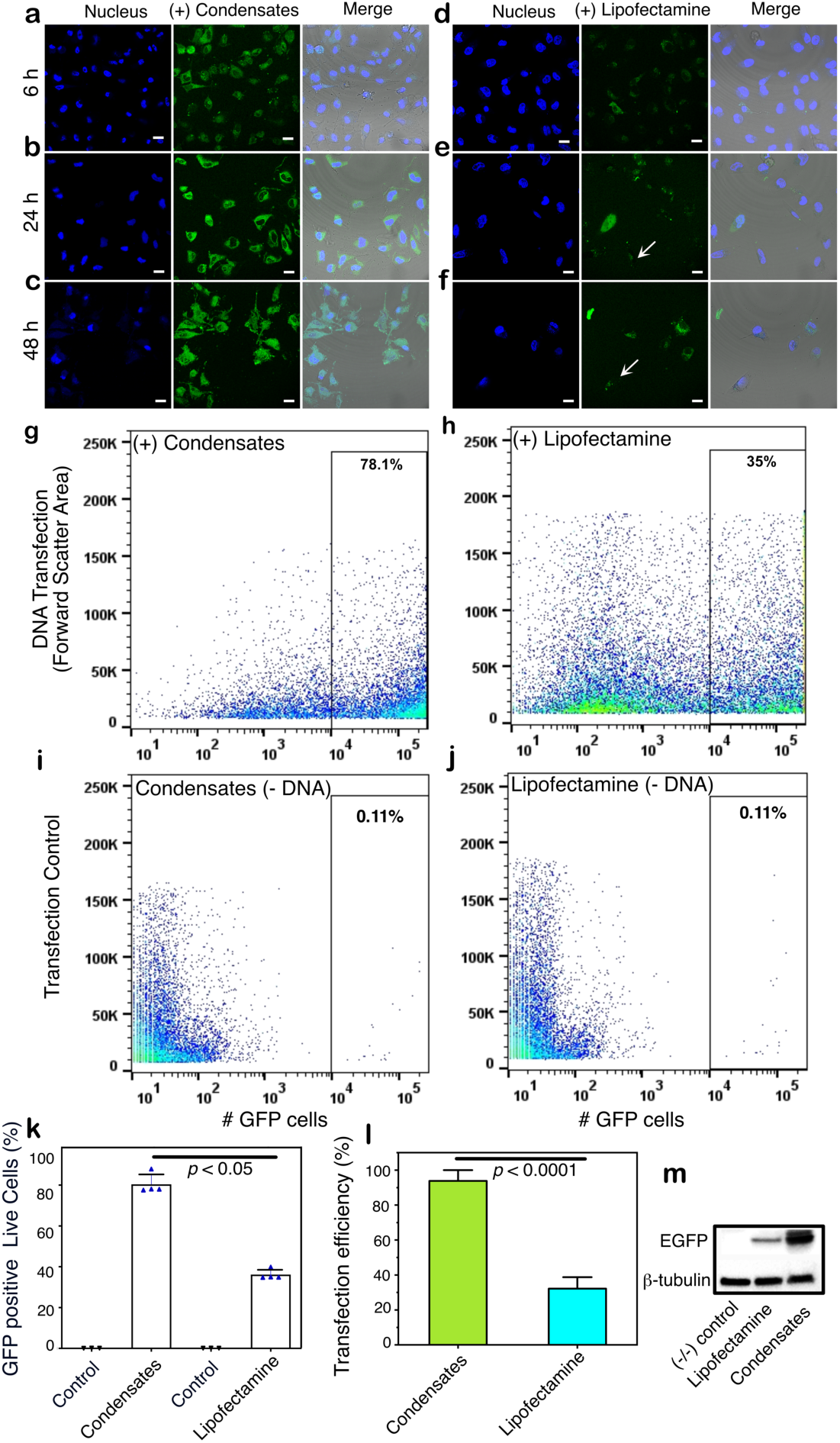
Efficiency of DNA delivery assessed by protein expression. Confocal microscopy images of HeLa cells showing the CMV-EGFP expression (green) upon transfection of its DNA by condensates (a–c) or Lipofectamine3000^®^ (d–f) after different incubation times as indicated (6, 24, and 48 h). Nuclei were stained with Hoechst33258^®^ (blue). The merged images include a brightfield overlay. The arrows indicate dead cells. Scale bars are 20 μm. (g-h) Live (APC^-^) GFP^+^ cells were analyzed via flow cytometry. Representative dot plots (GFP^+^ vs. FSC-A) display GFP expressions in live cells (APC^-^; corrected for dead cells), higher with condensates (78.1%) (g) than Lipofectamine (35.0%) (h). (i-j) Controls included cells treated with transfection reagents alone without the cargo. (k) Quantification of GFP-positive live cells (GFP^+^/ APC^-^); *n* = 3. (l) Quantitation of protein expression as transfection efficiency by condensates and Lipofectamine (see Methods). Statistical analysis was conducted using one-way (for k) and two-way (for l) ANOVA followed by Tukey’s test. Statistical significance is indicated. (m) Protein expression levels were assessed by western blot analyses of cell lysates.

Furthermore, many dead cells were observed with the Lipofectamine sample (arrows; Figure 7e and 7f) that were not present in cells transfected with PSP-2 condensates. Flow cytometry analysis of the live cells showed a significantly higher number of cells expressing GFP with condensate-transfection than the Lipofectamine-transfected ones (Figures 7g and 7h). Control samples without DNA showed a baseline of ∼11% cells (Figures 7i and 7j). Quantifying GFP expression in live cells showed a difference of 78.1% and 35.0% for condensates and Lipofectamine, respectively (Figure 7k). We also quantified transfection efficiencies based on the ratio of transfected cells to the total number of cells, which showed a similar difference in the transfection efficiencies with ∼88% for the condensates and ∼30% for Lipofectamine (Figure 7l). Quantitation of protein expression levels by western blot analysis also demonstrated a similar difference between the two transfection methods based on band intensities (Figure 7m). More importantly, cell death was minimal with the condensates compared to the Lipofectamine system. These data illustrate the efficiency of PSP-2 condensates to transfect HeLa cells with DNA to the cytosol, which eventually reaches the nucleus to undergo transcription.

We argue that if high encapsulation and transfection efficiency can be achieved with DNA using PSP-2 condensates, then mRNA could also be delivered with the same or better efficiency for two reasons: 1) PSP-2 is known to complex-coacervate with polyA-RNA, and 2) unlike DNA, mRNA has only to reach the cytosol (and the ribosomes in rough ER) to be effectively translated. To test this hypothesis, we used an mRNA transcript of GFP as cargo in PSP-2 condensates. We transfected HeLa cells with 1 µg of GFP mRNA through PSP-2 condensates. Within six hours of transfection, we observed bright green fluorescence in the cytoplasm of almost all the cells, confirming efficient transfection and expression of GFP (Figure 8a). The expression of GFP improved over the next 24 and 48 hours based on the fluorescence intensities in the cytoplasm (Figure 8b and 8c). Strikingly, almost all the cells were transfected by the condensates. In stark contrast, mRNA transfected with Lipofectamine3000^®^ showed good expression of the protein, but only a few cells were transfected during the same passage of time (Figure 8d-f). Flow cytometry analysis of the live cells showed a significantly higher number of cells expressing GFP with condensate- transfection than the Lipofectamine-transfected ones (Figures 8g and 8h), similar to DNA-transfection. Control samples without mRNA showed a baseline of ∼11% cells (Figures 7i and 7j). Quantifying GFP expression in live cells showed a difference of 75.3% and 28.8% for condensates and Lipofectamine, respectively (Figure 8i). We also quantified efficiencies based on the ratio of transfected cells to the total number of cells, which showed a similar difference in the transfection efficiencies with ∼80% for the condensates and ∼30% for Lipofectamine (Figure 8j). Quantitation of protein expression levels by western blot analysis also demonstrated a similar difference between the two transfection methods based on band intensities (Figure 8k).

**Figure 8.**
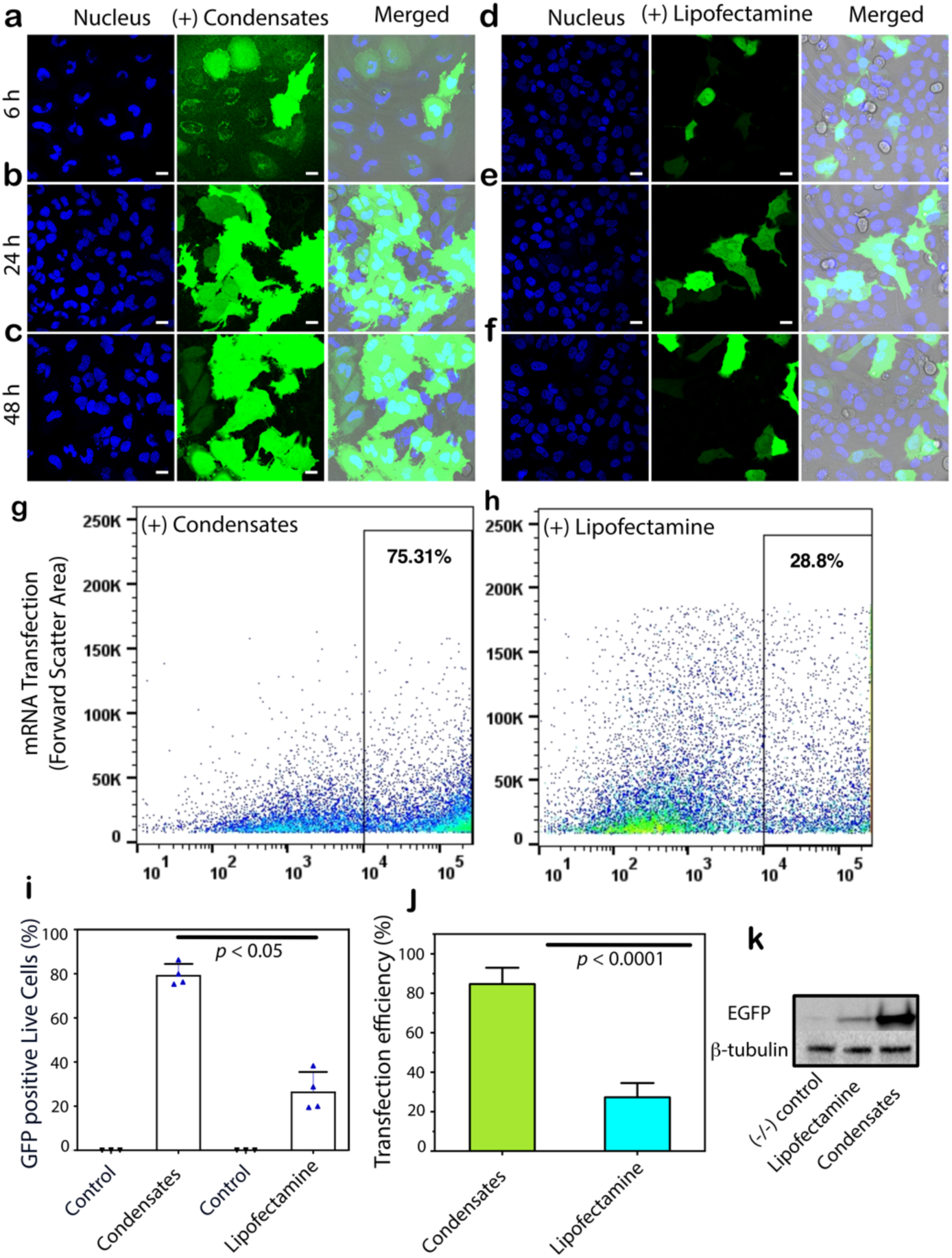
Efficiency of mRNA delivery assessed by protein expression. Confocal microscopy images of HeLa cells showing the GFP expression (green) upon transfection of its mRNA by PSP-2 condensates (a–c) or Lipofectamine3000^®^ (d–f) after different incubation times as indicated (6, 24, and 48 h). Nuclei were stained with Hoechst33258^®^ (blue). The merged images include a brightfield overlay. Scale bars are 20 μm. (g-h) Live (APC^-^) GFP^+^ cells were analyzed via flow cytometry. Representative dot plots (GFP^+^ vs. FSC-A) display GFP expressions in live cells (APC^-^; corrected for dead cells), higher with condensates (75.3%) (g) than Lipofectamine (28.8%) (h). (i) Quantification of GFP-positive live cells (GFP^+^/ APC^-^). *n* = 3. (j) Quantitation of protein expression as transfection efficiency by condensates and Lipofectamine (see Methods). Statistical analysis was conducted using one-way (for i) and two-way (for j) ANOVA followed by Tukey’s test. Statistical significance is indicated. (k) Protein expression levels were assessed by western blot of the cell lysates.

## CONCLUSIONS

There is an increasing demand for environment-sensing, efficient vehicles for the cellular delivery of therapeutics and molecular cargo. In our previous study, we demonstrated the design of phase-separating peptides (PSPs) containing cysteines that form stable and reversible condensates under redox flux^31^. We also showed that these condensates can partition small molecular cargo within them. Here, we have demonstrated the ability of PSP-2 condensates to effectively transport and deliver small and macromolecular cargo to the cytoplasm of HeLa cells. Our design principle for cargo delivery is based on the highly reducing environment of the cytoplasm^33^ where PSP-2 condensates, upon crossing the membrane, would spontaneously dissolve, releasing the cargo. The condensates’ precise cell entry mechanism remains unclear, and our ongoing investigations will uncover this in the future. However, several reports have already shown that condensates penetrate the membrane bilayer in mammalian cells and could do so through a mechanism without involving endosomal or membrane fusion^34–37^. Our initial investigations on the internalized condensates also support this contention and did not observe lipid membranes engulfing the condensates, which would indicate endocytosis (Figure S7a-c). We hypothesize the mechanism of internalization occurs as follows (schematically depicted in Figure 9). The positively charged PSP-2 condensates facilitate interaction and adhere to the negatively charged membranes on the extracellular side (Figure S6), as shown with other condensate systems^38^. Due to their interfacial tension, the condensates could bend and distort the membranes, as some are known to do to surfaces ^39–41^. Such deformations could allow the condensates to diffuse through the membrane bilayer passively. Once the condensates reach the cytosol, disulfide cross-links are reduced to thiols, dissolving the condensates to release the cargo. Although most PSP-2 condensates dissolve in the cytoplasm, some undissolved droplets are also observed (Figure 5). This partial condensate dissolution could indicate inhomogeneity in cytoplasmic redox fluxes. Although the cytoplasm was largely reducing, some localized oxidized zones were observed, as confirmed by the data on oxidizing environment sensing dye (Figure S7g-k). The cargo is also not transported to the lysosomes (Figure S7d-f). Such redox-based cargo delivery has also been used in pH-responsive positively charged peptides conjugated with a self-immolative disulfide moiety (HB*pep*-SR)^35–37^. Such peptide coacervates form stable interaction hubs in cells^42,43^. Irrespective of the cell permeation mechanism, our data unequivocally demonstrate that the cargo-laden condensates cross the membrane, dissolve, and efficiently deliver the encapsulated cargo to the cellular cytoplasm in an energy-independent manner.

**Figure 9.**
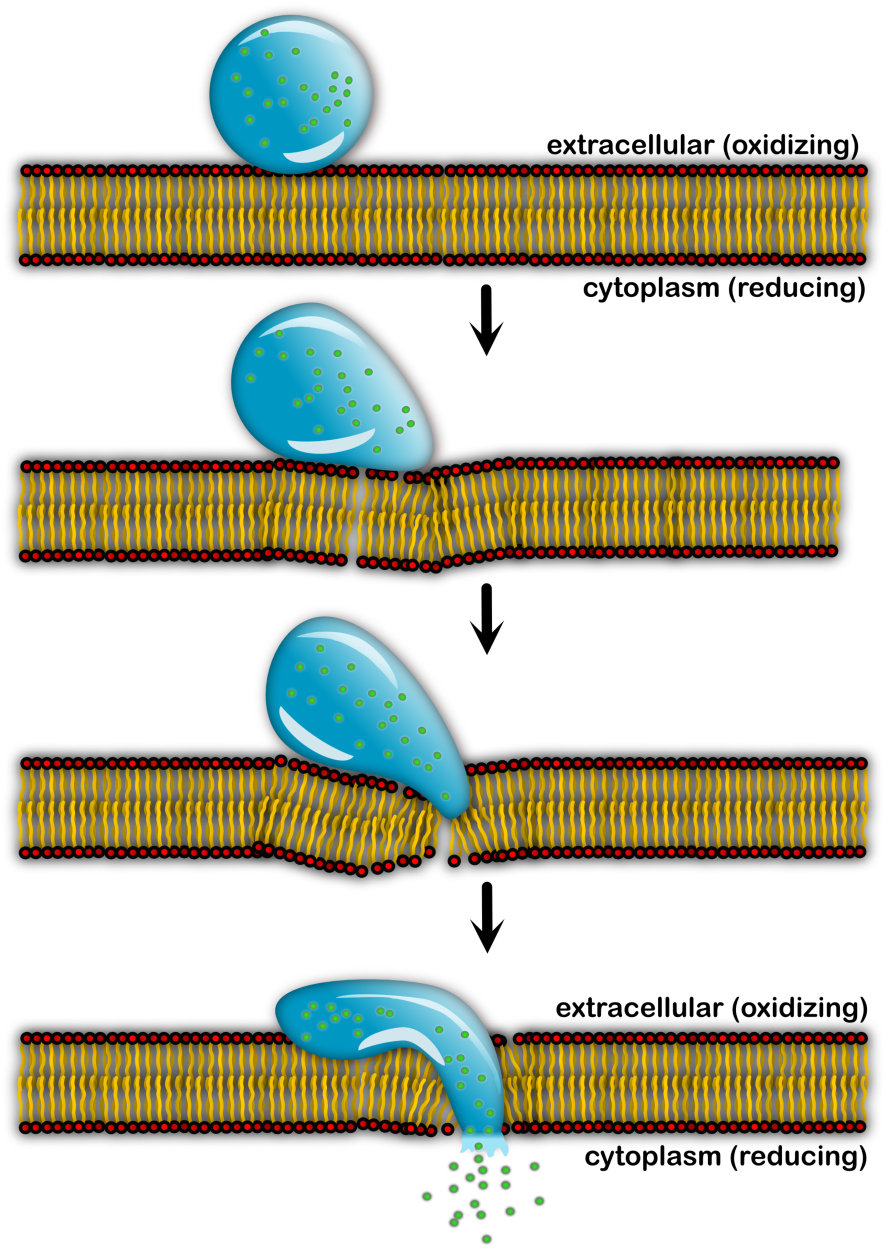
Hypothetical mechanism of cell permeation by PSP-2 condensates. The condensate adheres to the membrane surface on the extracellular side, diffuses through the membrane, and dissolves in the reducing cytoplasm to unload the cargo (green balls).

As novel delivery vehicles, our PSP condensates also offer many advantages over others, which are underscored by their tunability and customizability. PSP-2 condensates are large (∼5 μm on average) with a pore size of ∼100 nm diameter^31^ and can, therefore, accommodate various cargoes, from small molecules to biomacromolecules. As demonstrated here, PSP-2 condensates carry and efficiently deliver six different cargoes that vary in size and chemistry (Figure 1). Comparison between PSP-2 condensates and commercial lipid-based nanoparticle systems such as Lipofectamine showed that the condensates are significantly better in transfection efficiency for DNA and mRNA. Furthermore, PSP-2 is benign to HeLa cells as opposed to significant cell death caused by Lipofectamine, perhaps due to the cationic lipids in the system. The transfection efficiencies of the condensates can be further improved by modulating the encapsulation efficiencies that measure the partitioning of cargo depending on the nature of the cargo. In principle, PSP sequences can also be customized to cater to specific cell types, while redox sensitivity and pore sizes can be tuned by increasing the number of cysteines. The peptide sequences can also be modified to minimize potential cellular toxicity or immunogenicity without compromising the condensate-forming ability.

Many such customizable parameters make PSP condensate design promising to be used as versatile cell delivery vehicles.

## METHODS

### Materials

Rink Amide Protide Resin, Fmoc protected amino acids, and ethyl cyanoglyoxlate- 2oxime (Oxyma) were purchased from CEM peptides. Dichloromethane (DCM), diethyl ether, trifluoroacetic acid (TFA), N-dimethylformamide (DMF), acetonitrile, diisopropylcarbodiimide (DIC), triisopropylsilane (TIS), ethane-1,2-dithiol (EDT), and all other solvents were purchased from ThermoFisher Scientific or Sigma Aldrich at the highest purity. FITC and Fluorescein dyes were purchased from Thermo Scientific, mRNA of GFP was purchased from GeneScript^®^, and DNA plasmid of EGFP (pRSET his-eGFP) was purchased from Addgene^®^.

### Peptide synthesis

PSP-2 was synthesized as previously reported ^1^. Briefly, synthesis was carried out on a Liberty Blue 2.0 automated peptide synthesizer (CEM) through standard 9- fluorenyl methoxycarbonyl (Fmoc)-based solid phase peptide synthesis. Peptide synthesis was performed at 0.25 mmol scale using Rink Amide Protide Resin (0.65 mmol/g loading, 100- 200 mesh). Deprotection of Fmoc protecting groups was carried out using 20 v/v% piperidine in DMF. Each amino acid addition was carried out using Fmoc-protected amino acids (0.2 M), DIC (1M), and Oxyma (1 M) in DMF. After the final Fmoc deprotection, the resin beads were washed 3x using DCM. The peptide then underwent global deprotection and cleavage from the resin beads through gentle shaking in TFA/TIS/H_2_O/EDT (95: 2.5:2.5:2.5) cleavage cocktail for 4 hours at room temperature. Peptides were then precipitated in cold diethyl ether and collected via centrifugation. The peptide pellet was then resuspended in diethyl ether and chilled overnight at -20 °C. It was recentrifuged, and the diethyl ether supernatant was decanted from the peptide pellet, which in turn was allowed to air dry. Crude peptides were purified on a Prodigy preparative reverse-phase HPLC (CEM) with a water/acetonitrile gradient (containing 0.1% TFA). The mass and identity of the eluting fractions containing the desired PSP peptides were confirmed using electrospray ionization (ESI)- mass spectrometry (MS) on a Thermo Scientific Orbitrap Exploris™ 240.

### Preparation of PSP-2 condensates

Based on the *C_sat_* estimated in our previous study for PSP-2 self-coacervation, the lyophilized powder of the HPLC-purified peptide derivatives was dissolved in autoclaved, nanopure water, and the stock solution of ∼ 50-100 mM was stored at -20 °C until further use. Prior to the experiments, an aliquot of the peptide stock was taken to make a self-coacervate sample of 3.5 mM peptide concentration in 50 mM Tris, 2.5 M NaCl at pH 8.0. While producing complex coacervation, 200 µM of the peptide and 200 µg/mL Poly-A RNA in the presence of 150 mM NaCl (pH-7.4, 50 mM Tris buffer) was added.

### Turbidity assay

We employed turbidity as an indicator of LLPS, specifically for samples where liquid droplets were verified by optical microscopy. A BioTek Synergy H1 microplate reader was used to measure turbidity. Before each measurement, reactions were equilibrated for 10 minutes at 37 °C. All measurements were made at 37 °C. On Origin 8.5, data processing was carried out. At least three different datasets had their means determined.

### Reversibility and stability of PSP-2 condensates in cell culture medium

The reversible formation and dissolution of PSP-2 condensate in buffer and cell culture media after treatment of reducing and oxidizing agents was monitored using confocal microscopy. A cavity glass slide was filled with 50 µL of 3.5 mM PSP-2 in 50 mM Tris, 2.5 M NaCl, 0.5% FITC (as a reporter) at pH 8.0, air oxidized for 15 minutes before imaging. Following imaging, samples were treated with 5 mM DTT and re-imaged at reducing conditions. A 2% H_2_O_2_ (oxidizing agent) was added to the coacervates to observe the reformation of the condensates in confocal microscopy (Leica STELLARIS-DMI8 microscope). The stability and *C_sat_* of cargo- loaded condensates in Optimum^®^ media were determined by serial dilutions. After confirming the condensates, each cargo at 5 µM final concentration was added to the reactions imaged separately and incubated for 10 additional minutes. The condensates were then serially diluted in Optimum^®^ cell culture medium until no droplets were observed. The reversibility of cargo-free condensates in cell culture media was determined similar to the one in buffer but in Optimum^®^ media using both 5mM DTT and glutathione (GST) as reducing agents.

### Generation of CMV-EGFP DNA fragment

The pEG vector containing CMV promoter, EGFP ORF, and bGH (bovine growth hormone) poly(A) signal sequence (1731 bp) was constructed by in vivo cloning technique^44^. The vector backbone (F1, 5345 bp) containing Ori, Kan, CMV, and T7 promoters, and bGH poly(A) signal sequence was generated by PCR from the plasmid pcDNA3 Kan using primers TAAACCCGCTGATCAGCCT (BKf) and CATGGTGGCAAGCTTAAGTTTAAACGCTAGC (BKr). EGFP ORF (F2, 760 bp) was produced from pEGFP-N1 (Clontech) by PCR using CTTAAGCTTGCCACCATGGTGAGCAAGG (EGf) and CAGTCGAGGCTGATCAGCGGGTTTATTACTTGTACAGCTCGTC (Egr). The template DNA was removed by digestion with DpnI. The two DNA fragments were mixed at 3:1 ratio of F1:F2. One microliter of the mixture was used to construct plasmid pEG by in vivo cloning ^44^. The DNA fragment CMV-T7-EGFP-bGH poly(A) (1731 bp) was generated from pEG by PCR using primers CGATGTACGGGCCAGATATACGCGTTG (CMVf) and TcGATAAGATACATTGATGAGTCCCCAGCTGGTTCTTTCC (bGHr). The DNA fragment was EtOH-precipitated and dissolved in 100 μL water to make 10-fold concentrated DNA (relative to PCR reaction) for use in the experiment.

### Coating glass slides and coverslips

Coverslips (Fisher brand, microscope cover glass, 22x22 mm) and microscopic single cavity glass slides (GSC, Cat no: 4-13057-DZ-12) were sonicated for 15 minutes with 70% ethanol and air dried for 15 minutes. They were then immersed in 20% Tween20 solution for 30 minutes, following which they were rinsed eight to ten times with Milli-Q water to remove any remaining coating solution. Glass slides and coverslips were then dried overnight at 37 °C and wrapped in Lens paper until further use.

### Confocal microscopy and FRAP analysis

Confocal microscopic images of PSP-2 condensates (self- and complex-coacervates) on coated cavity glass slides (50 µL of reaction volume were added to cavity slide) were taken using a Leica STELLARIS-DMI8 microscope set to 63X magnification. Droplets were given a few minutes to settle in each response before being imaged. After droplets formed, 0.5% FITC was added to the reaction for imaging. The fluorescence recovery after photobleaching (FRAP) technique was used to study the internal dynamics of the self and complex condensates. The liquid droplets were photo-bleached by exposing them to a laser with an intensity of about 90% for five seconds. For sixty seconds, the fluorescence was seen to recover. Origin 8.5 was used to normalize and visualize the fluorescence recovery kinetics against time.

### Cytotoxicity by lactose dehydrogenase (LDH) assay

HeLa cells were cultured in T25 flask to 90% confluence in an incubator set at 37 °C with 5% CO_2_. DMEM supplemented with 10% FBS and 1% penicillin-streptomycin was used to cultivate the cells. To enable cell adhesion, cells were then plated in a 96-well plate (1 × 10^4^ cells/mL, 100 μL volume per well) and incubated overnight at 37 °C and with 5% CO_2_. A 3.5 mM stock of peptide solution made in Tris buffer without salt and 3.5 mM of the peptide incubated with 2.5 molar NaCl to induce self-coacervation (pH-8.0, 50 mM Tris buffer). This was followed by dilutions with DMEM to achieve the final treatment concentration with cells. Each concentration was assessed in triplicate.

Triton X-100 served as a positive control for 100% cytotoxicity, or full LDH release, while nuclease-free water served as a spontaneous LDH control. Before collecting media to measure LDH release using the CyQUANT™ LDH Cytotoxicity Assay Kit in accordance with the manufacturer’s instructions, plates were incubated for 24 hours at 37 °C and 5% CO_2_.

### Encapsulation efficiency

Peptide solutions were oxidized and diluted as described above. To each solution, 5 µM of cargo and 100 ng of DNA were added to droplets containing reactions and allowed to incubate for 10 minutes. Standard solutions for each cargo and DNA of equivalent concentration were prepared in buffer and their UV-vis absorbances were recorded. The reactions (droplets containing) were centrifuged at 3000 xg for 10 minutes. Of the 100 µL volume, 40 µL was drawn as the supernatant, and the UV-vis absorbance was measured. The encapsulation efficiency of the cargo and DNA were calculated by following.

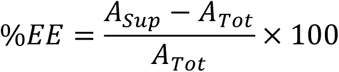

### Cell culture

Wild-type HeLa cell lines were a kind gift from Dr. Hao Xu from the University of Southern Mississippi. HeLa Cells were cultured in Dulbecco’s modified Eagle’s medium (DMEM) (Gibco) supplemented with 10% fetal bovine serum and maintained at 37 °C in a humidified incubator with 5% CO_2_. The subculture started by detaching the cells with trypsin treatment, followed by centrifugation (3000×*g*, 5 min) to collect the cells. Then, the pellets were resuspended with fresh media for subculture experiments.

### Cargo delivery

To perform cargo deliveries, 2 × 10^5^ cells were seeded in 1 mL media (12 wells plate). After incubating at 37°C in a humidified incubator with 5.5% CO_2_ overnight, the medium was replaced with 900 μL of Opti-MEM, then 100 μL of freshly prepared cargo encapsulated condensates (around 350 µM) were added into the Opti-MEM (Condensates were formed by incubating 3.5 mM peptide in 50 mM Tris buffer at pH 8.0, 2.5 M NaCl, then 5 µM of each cargo was added to the reaction and kept for 10 minutes). After 4 h of incubation, the cargo-containing medium was removed, and the cells were washed with PBS twice before being cultured in 1 mL of fresh DMEM. Then treated live cells were imaged at different time points using confocal microscopy.

### Transfection

EGFP-encoding DNA and mRNA of EGFP were used to evaluate the transfection efficiency of the PSP-2 self coacervates. Before transfection, Hela cells were incubated in 12 well plates with a density of 2 × 10^5^ cells per well for 12 hours. The medium was replaced with 900 μL of Opti-MEM, followed by the addition of 100 μL of freshly prepared EGFP DNA and mRNA-loaded coacervate suspensions (around 350 µM). For transfection ∼ 1 μg of DNA and mRNA were transfected. After 4h of incubation, the medium was removed, and the cells were washed with PBS twice before adding 1 mL of medium (DMEM, 10% FBS, antibiotics). The transfection was conducted for four hours of uptake, and protein expression was checked by imaging under a confocal microscope at 6, 24, and 48 hours and testing the transfection efficiency by western blot and ImageJ analysis.

### Transfection efficiency and Western blot analysis

Transfection efficiency was calculated by analyzing confocal images using ImageJ (For each sample, five images were analyzed, around 200 cells). First, the total number of cells was counted based on nuclear staining, and then transfected cells were calculated with GFP expression. Then, the % transfection efficiency was calculated by dividing the number of cells with detectable fluorescence (GFP) by the total number of cells and multiplying by 100. For western blot, transfected cells were lysed by Thermo Scientific NE-PER Nuclear and Cytoplasmic Extraction Kit after 24 hours. Aliquots of the fractionated samples were then separately mixed in the 4x Laemmli sample buffer and loaded onto SDS-PAGE Biorad Mini-PROTEAN^®^ 4-20% precast gel. Gels were then transferred onto a 0.45 μM Amersham Protran Premium nitrocellulose membrane (GE Life Sciences), and the blot was boiled in 1X PBS for one minute. Blot was then incubated for one hour in the blocking buffer (5% non-fat milk, 0.1% Tween^®^−20 in 1X PBS) and followed by anti-GFP primary antibodies (ProteinTech) overnight at 4°C. The following morning, the blot was incubated for an hour at room temperature after washing the blot 3 times with Tris buffer saline (TBS) with 20 % tween and horseradish peroxidase-conjugated anti-rabbit secondary antibodies. The blots were then imaged after treating them with ECL reagent on a GelDoc molecular imager (Bio-Rad).

### Flow Cytometry Analysis

HeLa cells were transfected with plasmid DNA or mRNA encoding GFP using either condensates or Lipofectamine™ 3000 (Invitrogen). HeLa cells with transfection reagents alone (condensates or Lipofectamine) were included as transfection controls. Following transfection, cells were detached by trypsinization and stained with LIVE/DEAD^TM^ Fixable Dead Cell Stain (APC-conjugated; Thermo Scientific) to discriminate live and dead cell populations. After staining, cells were incubated, washed, centrifuged, and resuspended in PBS for flow cytometry. All samples were analyzed using a BD FACSMelody^TM^ cell sorter, and data were processed using FlowJo software (v10.9.0). Quantitative analysis of GFP-positive live cells (FITC^+^/APC^-^) was performed, and bar graphs representing averaged results were generated using GraphPad Prism (v10.2.3).

### Image Processing and Analysis

A custom pipeline designed in R (version 4.1.2) and FIJI (Fiji Is Just ImageJ) version 1.54f was used to handle and analyze confocal microscopy images in order to determine the size distribution of the phase-separated droplets across time (both complex and self coacervates). The photos were pre-processed in ImageJ using a despeckle filter to eliminate noise and Huang’s auto-thresholding approach for binarization ^3^. Droplet segmentation was done using the watershed algorithm and morphological processes (erosion and dilatation) ^4^. After removing droplets that touched the frame’s edge, the segmented droplets were examined, and their area, mean intensity, perimeter, and shape descriptors were measured and exported as CSV files. In R, the CSV files from ImageJ were processed using custom scripts. The tidyverse, ggsci, and scales packages were utilized for data wrangling, visualization, and statistical analysis ^5–6^ . The droplet counts and size distributions were summarized for each combination of peptide, time point, and replicates. Boxplots were generated to visualize each peptide’s droplet size distribution across time points. The image analysis pipeline assumed that the phase-separated droplets were spherical and did not account for potential deviations from this shape. The segmentation process may have also introduced errors for closely spaced or overlapping droplets, leading to potential undercounting or inaccurate size measurements.

## AUTHOR INFORMATION

Conceptualization – VR; intellectual inputs – VR, MM, WSS, TDC; biophysical and cell biology experiments, data interpretation – MM; peptide synthesis – AMD, WSS, TDC; CMV EGFB design – FH; SUK, SS, FB – Flow cytometry analysis; data processing – MM; manuscript writing and editing – VR, MM, WSS, TDC, FB, FH.

## ACKNOWLEDGEMENTS

The authors thank the National Science Foundation (BMAT 2208349 to VR and TDC) and Mississippi SMART ACT Accelerator Initiative (VR) for their financial support. The authors also thank the National Center for Research Resources (5P20RR01647-11) and the National Institute of General Medical Sciences (8 P20GM103476-11) from the National Institutes of Health (NIH) for funding through INBRE to use their core facilities, especially confocal microscopy. Additionally, the authors acknowledge support from the National Science Foundation (2319932) for mass spectrometry instrumentation utilized in this study. TDC acknowledges funding support from the NIH and the National Institute of Biomedical Imaging and Bioengineering (NIBIB R21EB033533). TDC and AMD gratefully acknowledge support from the Arnold and Mabel Beckman Foundation. The authors also thank Ms. Azin Mirzazadeh for her help with cell culture experiments.

## TOC Figure

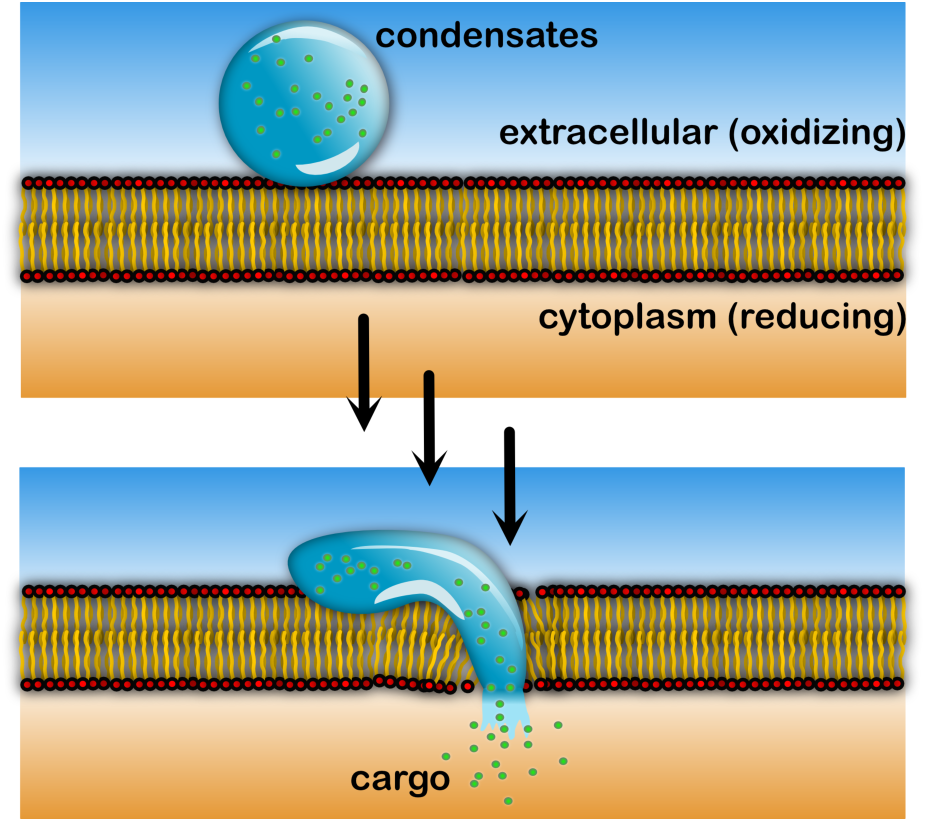

## SUPPLEMENTARY FIGURES

**Figure S1.**
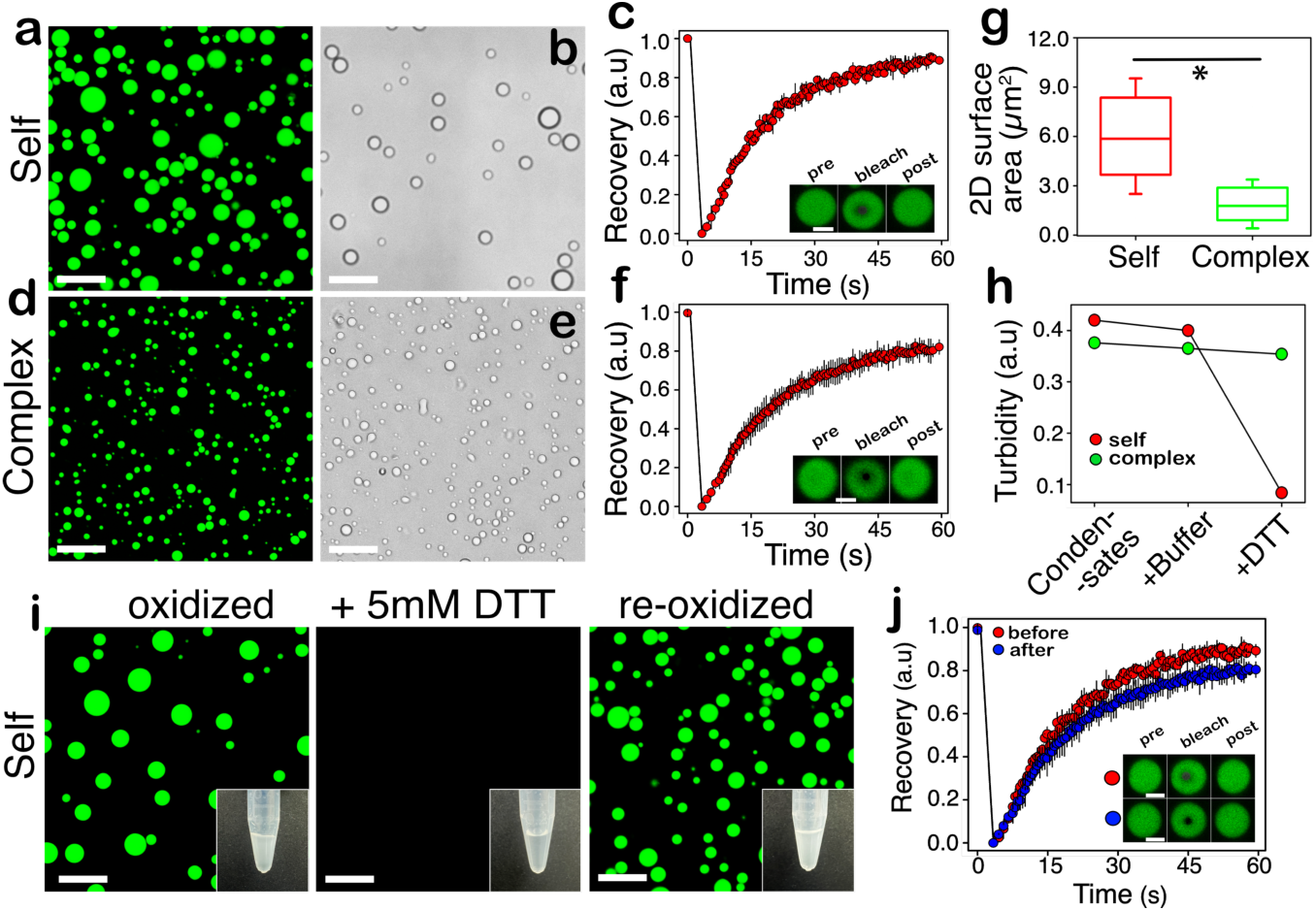
Characterization of PSP-2 condensates in buffer. (a) Confocal microscopy and (b) brightfield images of 3.5 mM PSP-2 peptide self-coacervates in 50 mM tris buffer with 2.5 M NaCl, pH 8.0 at 37 °C. (c) Confocal microscopy and (d) brightfield images of complex-coacervates containing 200 µM PSP-2 with 200 µg/mL PolyA-RNA in the same buffer but with 50 mM NaCl. For confocal visualization of the droplets, 1% FITC-labeled peptide was added to the reactions. (e, f) Fluorescence recovery after photobleaching (FRAP) analysis of condensates formed by self- and complex-coacervation, respectively, along with the regions of interest (ROIs; insets). (g) Sizes of the droplets were quantified based on the 2D surface area observed in the images using ImageJ software. * p<0.05. (h) The dissolution of condensates was demonstrated by turbidity; the reducing agent, DTT, was added to a final concentration of 5 mM. An identical volume of buffer without DTT was added as a negative control. (i) Confocal microscopy images of FITC-labeled PSP-2 peptide self-coacervates under respective LLPS conditions. The air-oxidized condensates were first dissolved using DTT and re-oxidized using hydrogen peroxide. The insets show Eppendorf tubes in their respective conditions. (j) FRAP analysis to assess the viscosity of the self-coacervates before and after reoxidation along with ROIs in the insets. The scale bar is 20 µm.

**Figure S2:**
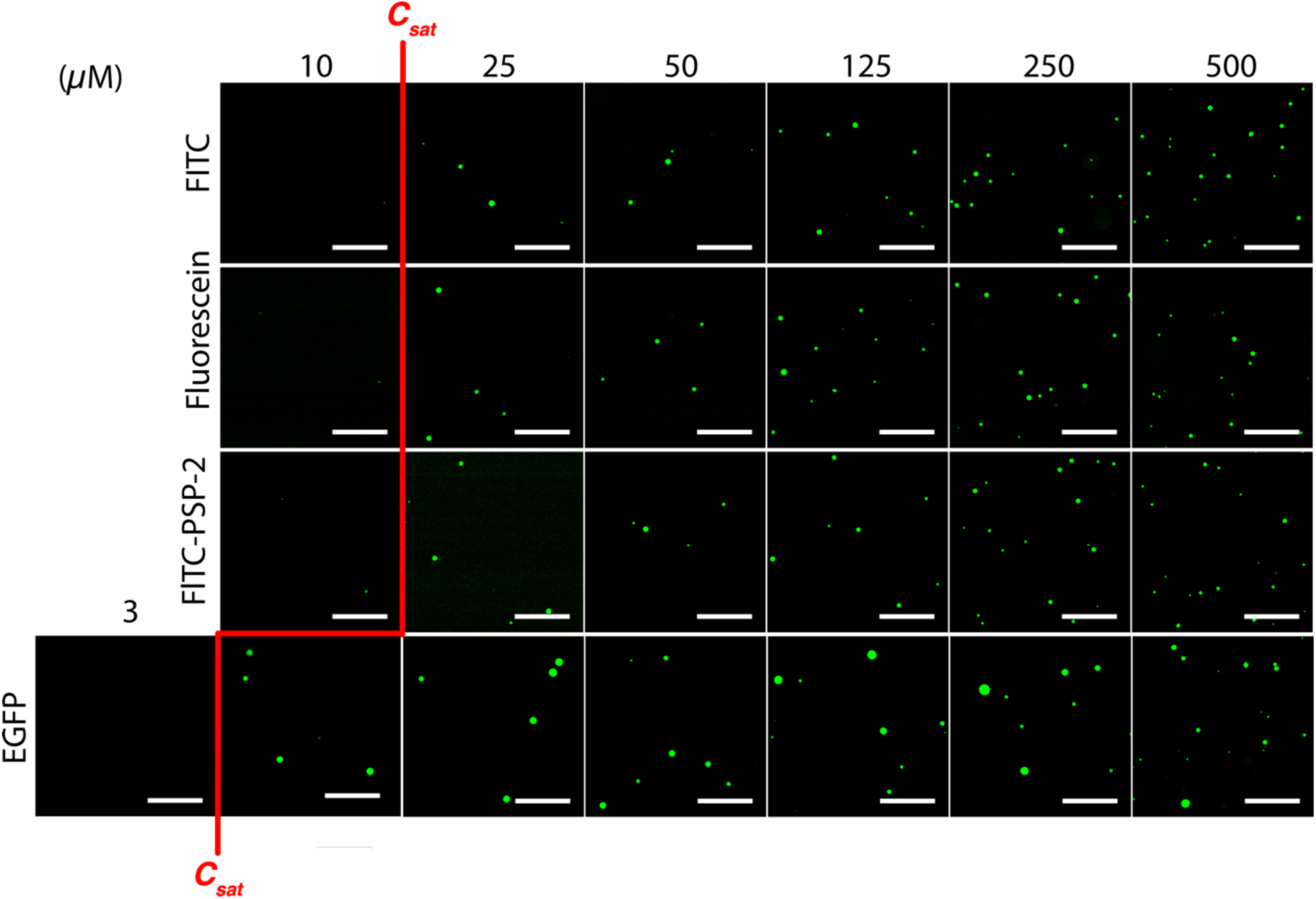
Stability of the different cargo-loaded droplets in the optimum medium at different time points. Confocal images of cargo-loaded PSP-2 self-coacervates in optimum media for FITC, fluorescein, FITC-labeled peptides, or EGFP protein at different dilution states (10 µMM,25 uM,50 uM,100 uM,250 uM and 500 uM). The scale bar is 20 µm.

**Figure S3:**
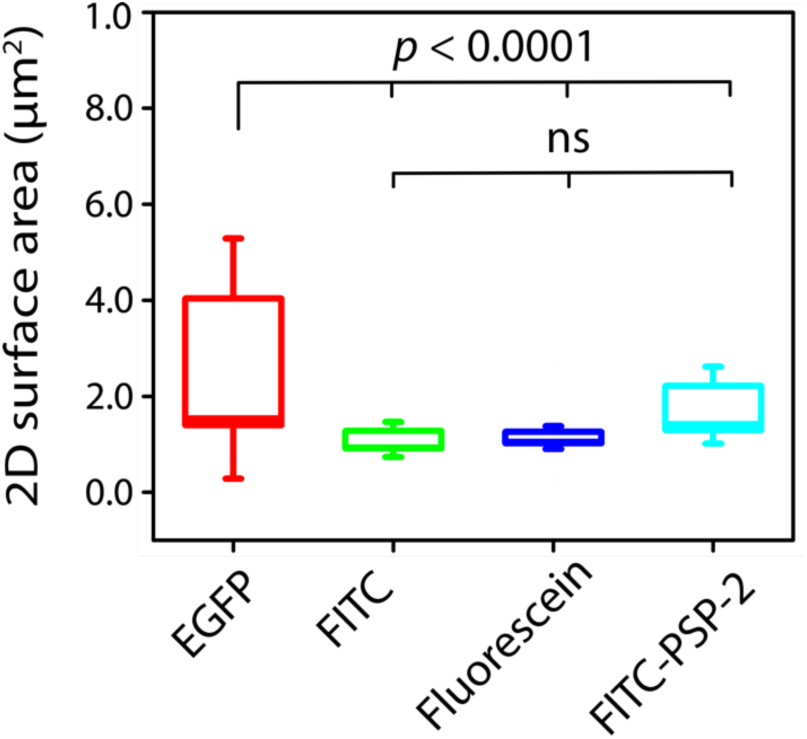
Size of PSP-2 condensates after loading cargo. Sizes of the cargo-loaded self-coacervates in Optimum media after 4 hours of incubation were quantified based on the 2D surface area observed in the images using ImageJ software for EGFP protein, FITC, fluorescein, and FITC-labeled peptides.

**Figure S4:**
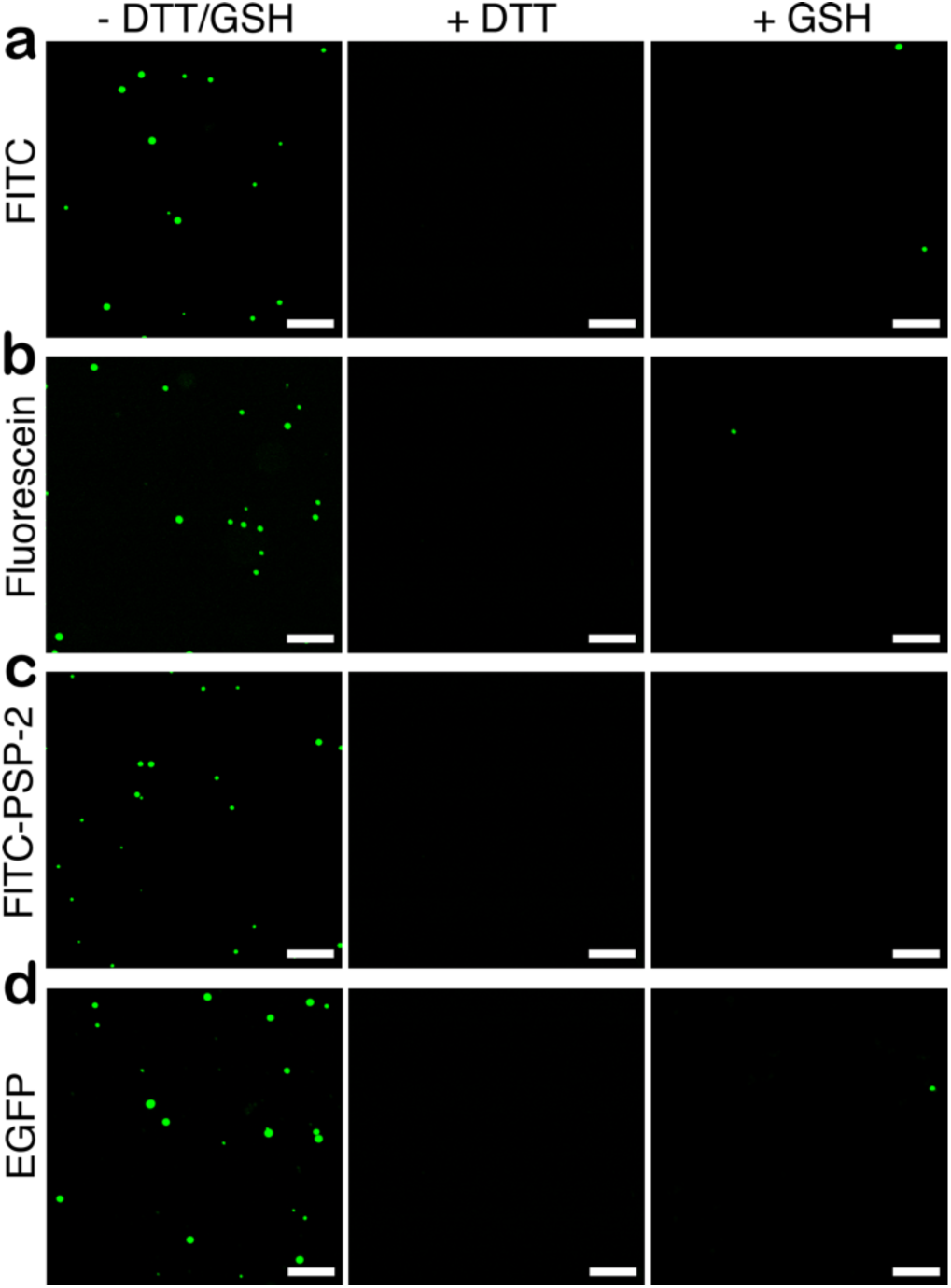
Cargo-laden PSP-2 condensates in Optimum media dissolve in reducing conditions. Confocal microscopy images of different cargo-loaded PSP-2 peptide self-coacervates for FITC (a), fluorescein (b), FITC-labeled peptides (c), and EGFP protein (d) under their respective LLPS conditions. Then, droplets were dissolved using 5mM DTT or GSH. Complete dissolution of condensates was observed after 4 hours incubation with GSH. The scale bar is 20 µm.

**Figure S5:**
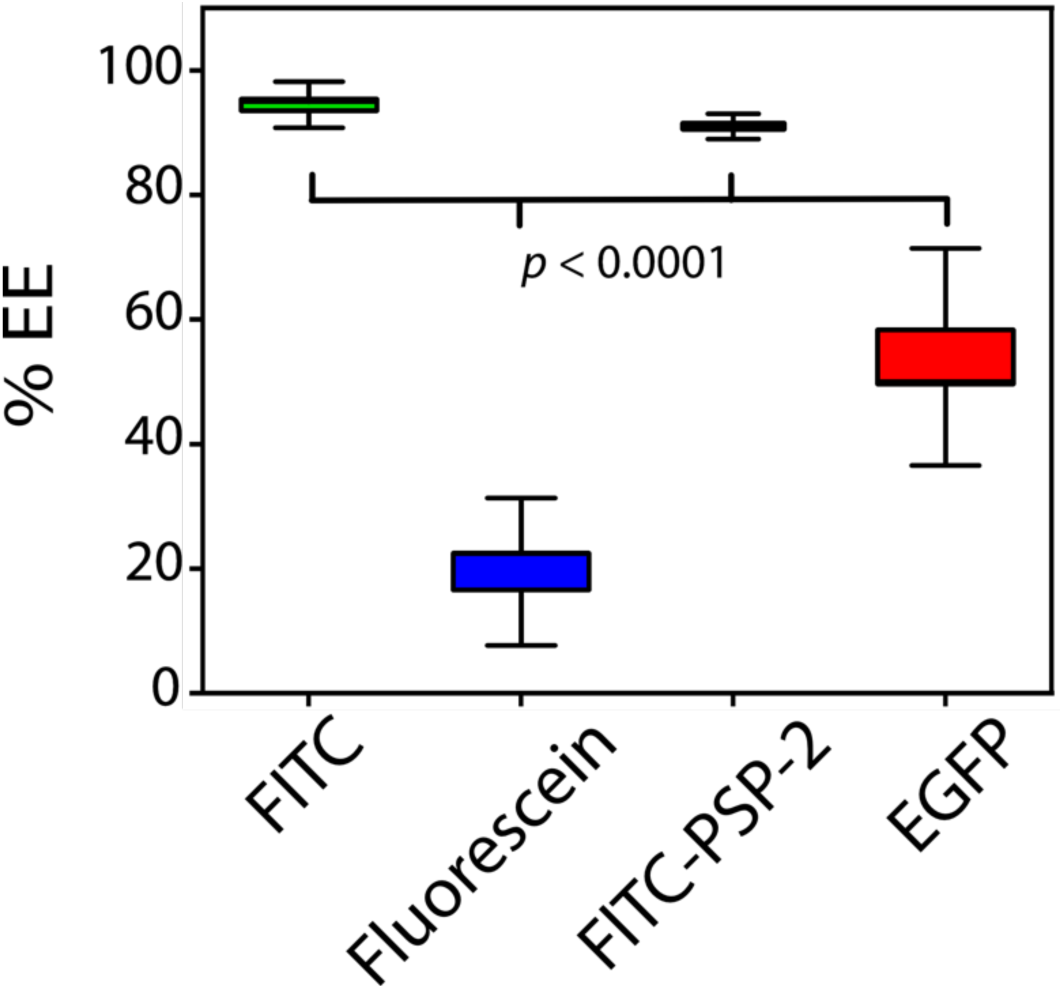
Encapsulation efficiencies of cargo in cell medium. The encapsulation efficiency (%) was assessed for different cargo in Optimum^®^ media. An average of three independent experiments are averaged. Two-way ANOVA between the data sets is shown (*P* < 0.0001).

**Figure S6:**
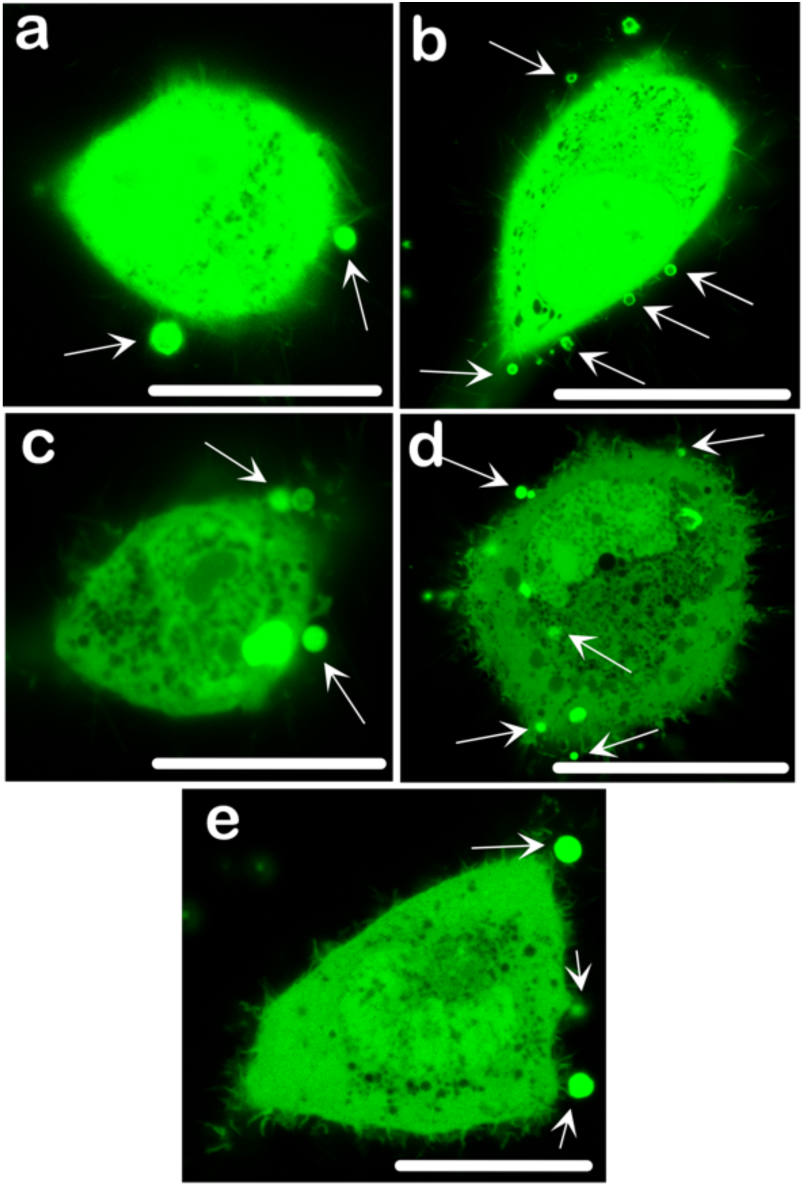
PSP-2 condensates have an affinity for membranes on the extracellular side. Confocal images of EGFP partitioning in HeLa cells show some condensates associated with membrane surfaces on the extracellular side, as indicated by white arrows.

**Figure S7:**
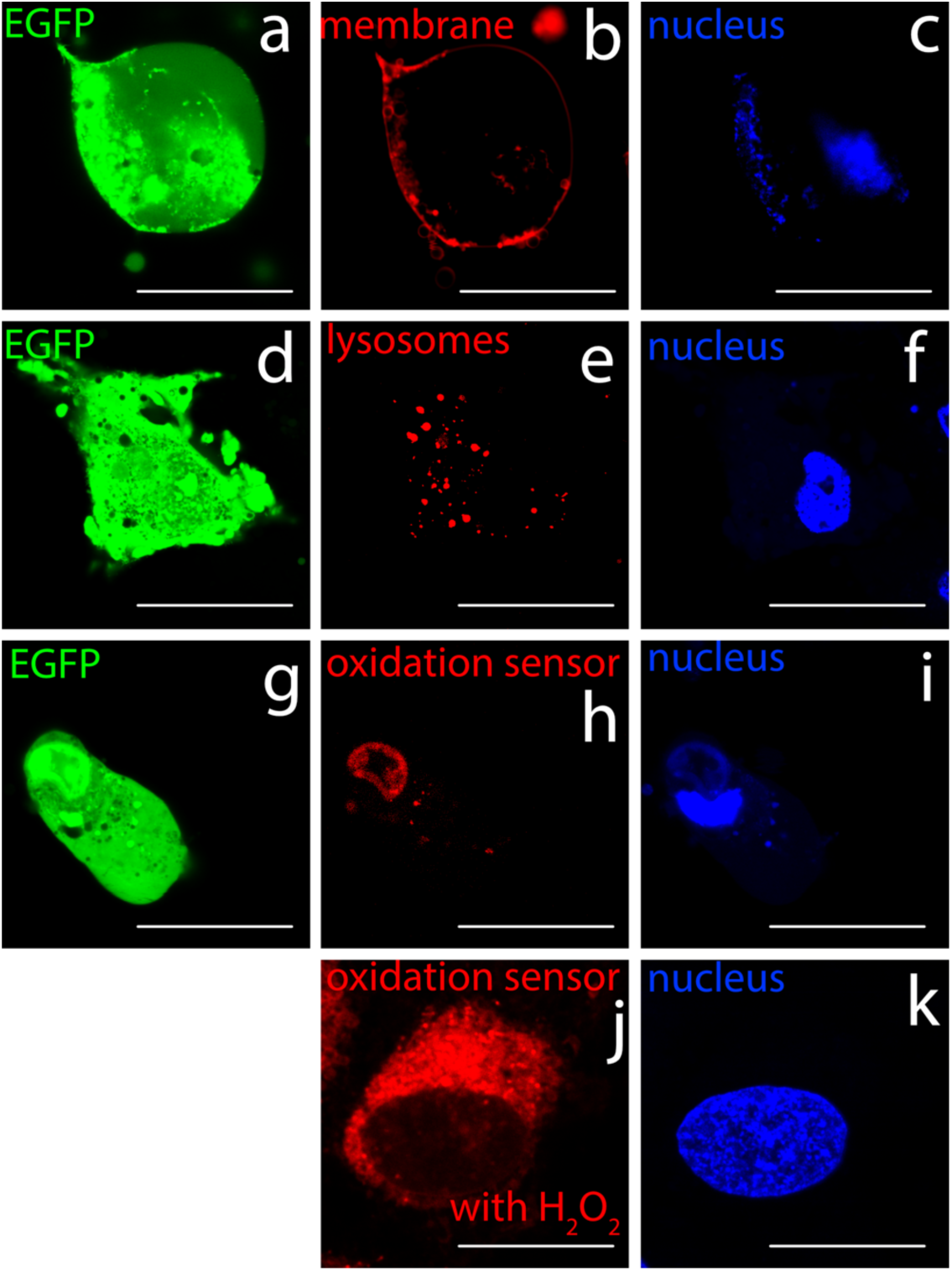
Cellular environment of condensates. Confocal images of EGFP partitioning in HeLa cells. In addition to EGFP (a, d, g) and nucleus (c, f, I, k), the cells were orthogonally stained for membranes (Cellbrite Fix 640 membrane Dye (Biotium)) (b), lysosomes (LysoView 650 (Biotium)) (e), and oxidation sensor (CellROX Deep Red Reagent (Invitrogen)) (j, j).

## Notes

### Competing Interest Statement

The authors have declared no competing interest.

